# Finnish gyrate atrophy mutation OAT;c.1205C>T leads to accumulation of intracellular GABA

**DOI:** 10.1101/2024.05.13.593857

**Authors:** Rocio Sartori-Maldonado, Kirmo Wartiovaara

## Abstract

Hyperornithinaemia with gyrate atrophy of choroid and retina (HOGA) is a recessive metabolic disease caused by dysfunction of the ornithine aminotransferase (OAT) gene, leading to ornithine accumulation and a complex metabolic imbalance. This causes retinal degeneration that ultimately evolve to blindness. However, the mechanisms of this degeneration remain unknown. Here, we have conducted untargeted metabolomic analysis in patient-derived induced pluripotent stem cells and their isogenic counterparts. Mutant cells show altered levels of ornithine-related metabolites, including low creatine, proline and glutamate, and elevated arginine and citrulline. The untargeted metabolomics approach revealed changes in the urea cycle and polyamine synthesis pathways with a significant intracellular accumulation of gamma-aminobutyric acid (GABA). Hence, we propose GABA as a key player in the disease pathogenicity, potentially affecting neuronal function in the eye.

## Introduction

Hyperornithinaemia with gyrate atrophy of choroid and retina (HOGA), also known as gyrate atrophy, is a childhood-onset recessive metabolic disease manifesting with nyctalopia, type-II muscle fibre degeneration, and retinochoroidal atrophy that progressively leads to blindness in adulthood^1–3^. The pathologic mutations behind this disease occur in ornithine aminotransferase (*OAT*) gene, with the Finnish-founder mutation OAT c.1205 T > C p.(Leu402Pro) leading the prevalence ranking and present in most of the Finnish patients ^4,5^.

The gene-homonymous protein (OAT) lays at a crossroad of metabolic pathways. Hence, the loss of its enzymatic activity leads to complex metabolic changes, mostly consequent to the accumulation of ornithine^5–7^. To off-set the effects of high ornithine concentrations, the most effective treatment to date involves dietary management with strict arginine restriction and supplementation with creatine and lysine^8–12^. In animal models, this diet completely prevents the retinal degeneration, while in humans it only delays the progression of the symptoms^11–13^.

A recent article reported a gene replacement strategy delivering a functional copy of *OAT* to the liver and showed improved function and structure of the murine retina. Another report from our group revealed an increase in the transcription of the gene in mutant cells^14^. Thus, the promise of the novel therapeutic approach still requires further research to confirm the genetic and molecular dynamics between the mutant and the replacement sequence, the efficiency in human patients and the actual impact on disease progression.

In Maldonado et al. 2022, we additionally introduced the first patient-derived induced pluripotent stem cell (iPSC) model for the disease, including the genetic correction of the cells, analysis of OAT expression and ornithine concentrations, and targeted metabolomics of some ornithine-related metabolites. Here, we have expanded that study to include intra– and extracellular untargeted metabolomics, aiming to widen our understanding on the disease mechanism. Our results showed complex dynamics on metabolites management and transport, with different changes inside of the cell than in the cell media. Among these changes, we found a striking intracellular accumulation of gamma-aminobutyric acid (GABA), which may explain some of the HOGA pathogenicity.

## Results

### Protein recovery

Previous reports suggested that the change c.1205 T>C in *OAT* destabilises the structure of the protein and leads to its degradation^7^. Accordingly, despite the two-fold increase in mRNA expression of the *OAT* gene in the mutant cells^14^, they lack the protein. The exact mechanism behind this event remains unexplored. Nevertheless, protein modelling with Uniprot (P04181, PDB ID: 7TA1) and AlphaFold 2.0 showed that the leucine (Leu) in position 402 locates at a crossroad between secondary protein structures, following a short loop and holding interaction with both beta sheet and alpha helix. (Fig.1). Its replacement by a proline (Pro), a conformationally constrained amino acid, likely introduces rigidity to the structure, disrupting the Leu-containing beta sheet. Furthermore, the loss of one hydrogen atom in the protein, consequent to Proline’s pyrrolidine ring, reduces the number of stabilising hydrogen bonds. Further experimental and computational experiments may shed light onto these events.

**Figure 1.**
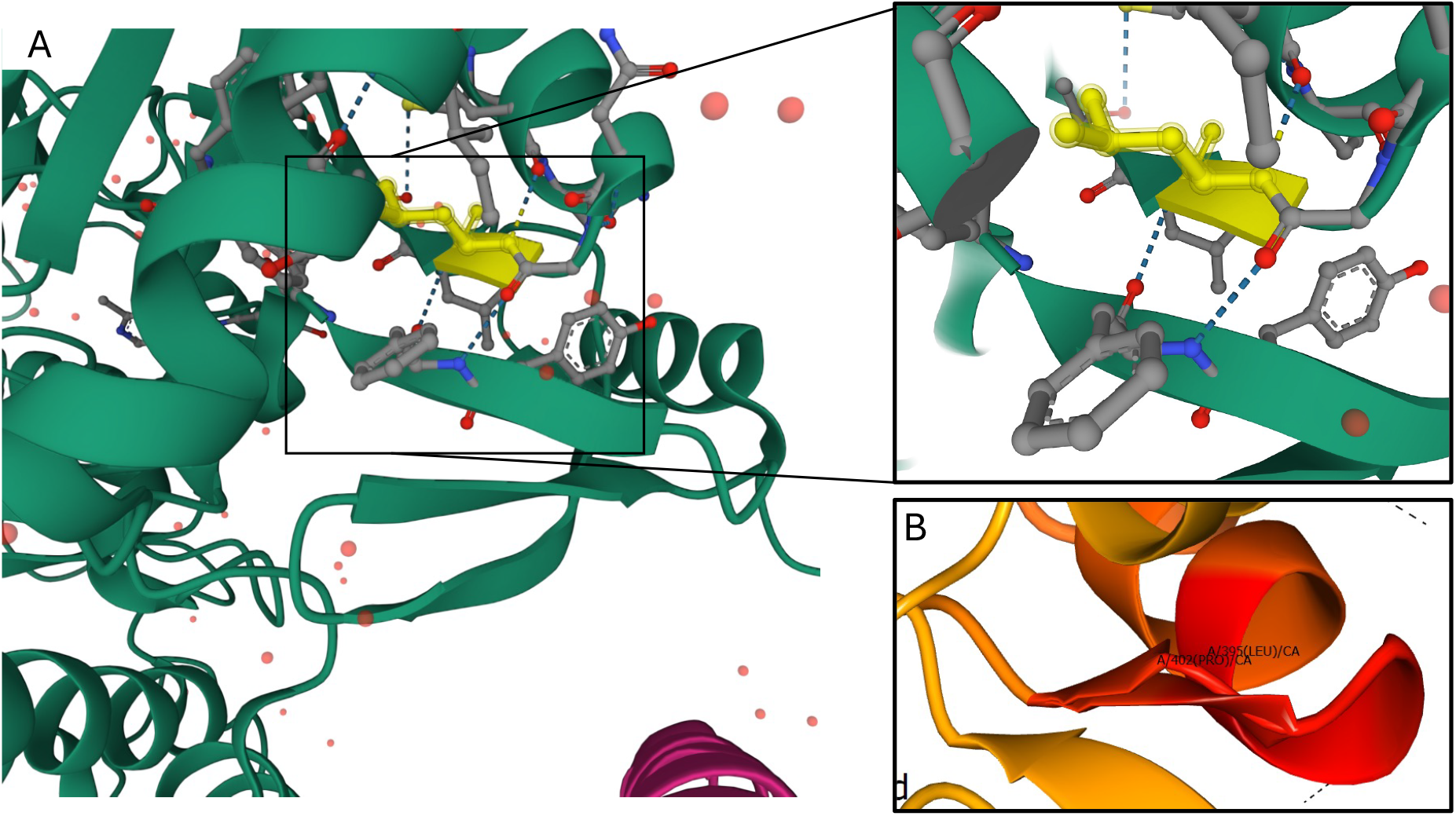
**A**. Model of protein P04181 (OAT) showing Leucine and the multiple hydrogen bonds it forms. **B**. Change in secondary structure in OAT:p.L402P (shorter B sheet), overlapped with the wildtype structure (longer B sheet).

### Restoration of metabolic homeostasis

After our initial reports on metabolic changes in mutant cells compared to CRISPR/Cas9 corrected cells, we sought to understand the wider metabolic landscape with exploratory untargeted metabolomics in both cellular media and cell lysates. The analysis identified some distinct metabolic dynamics in the cell, illustrating different strategies to maintain metabolic homeostasis upon high concentrations of ornithine. We initially selected some ornithine related metabolites, some of which present reported changes in the patients^3,15–18^. As expected, mutant cells show a significant increase in ornithine levels, as well as a marked decrease in proline, glutamine, and glutamate, directly related to the OAT dysfunction (Fig. 2A-D). Surprisingly, contrary to our previous reports, mutant cells present significantly higher arginine levels. Nevertheless, the concurrent increase in citrulline may suggest switches in the cellular strategies to deal with ornithine at the time of sampling. Thus, the untargeted metabolomic approach suggests that initially ornithine diverts through the urea cycle. Additionally, the sharp decrease of creatine and phosphocreatine in both media and lysates in mutant cells compared to their corrected counterparts indicates an active catabolic path from Arginine. Further enrichment analysis confirmed the upregulation of ornithine processing and related pathways, including the urea cycle and the tricarboxylic acid (TCA, also known as citric acid cycle) cycle, in both cell media and cell lysates (Fig. 2E,F, supplementary figure 1). Some enriched intracellular metabolites, such as cyclic GMP (cGMP) and GABA, supported by the upregulation of ornithine decarboxylase (ODC, Fig 2F), suggest the activation of the polyamine synthesis pathway. These metabolites, however, do not increase in the cell media. The striking high levels of intracellular GABA in c.1205T/T cells may indicate a lack of excretion of this metabolite, but further experiments should be conducted. Instead, the cell seems to transform this GABA into other metabolites, like succinate and glutamate, that later filter onto the cell medium. The imbalance of these and other metabolites inside the cell might explain the combination of enriched energetic pathways, as gluconeogenesis, glutamate metabolism, and the citric acid cycle. On the other hand, the enrichment in the culture media reflect the effect of ornithine handling, representing pathways more directly linked to ornithine and ornithine-related metabolites, such as the urea cycle, citric acid cycle, and arginine, proline, and glycine metabolism. Moreover, we see an upregulation of the cationic amino acid transporter (CAT1, Fig 2F), which mobilises ornithine through the plasma membrane, further suggesting active transport of metabolites in an out of the cell.

**Figure 2.**
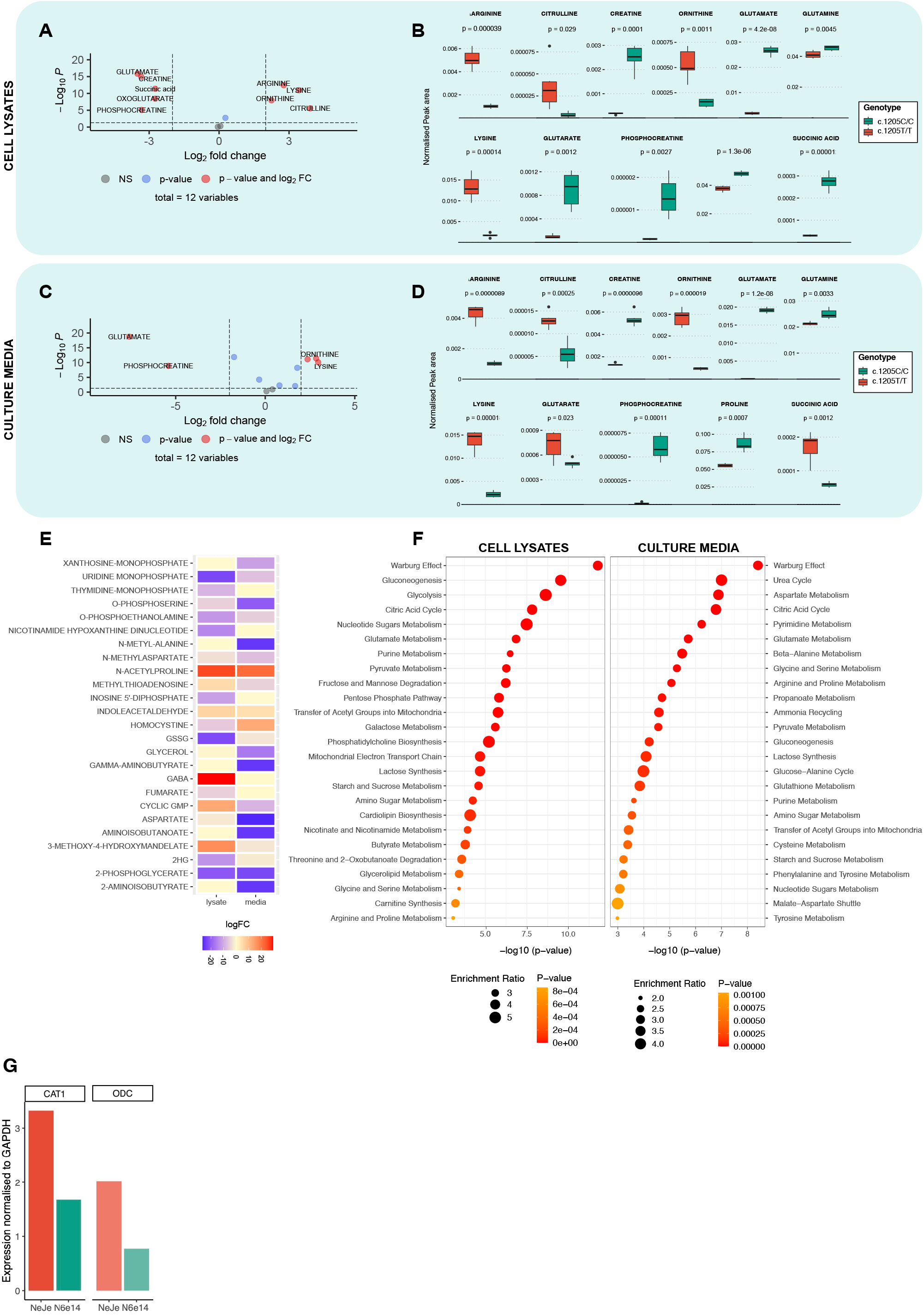
**A**. Volcano plot of depicting changes in targeted ornithine-related metabolites in cell lysates from patient iPSC, compared to the corrected cell lines. **B**. Levels of targeted metabolites in mutant and corrected cell lysates, normalised to total metabolites. **C**. Volcano plot of depicting changes in targeted ornithine-related metabolites in cell culture media from patient iPSC, compared to the corrected cell lines. **D**. Levels of targeted metabolites in mutant and corrected cell lysates, normalised to total metabolites. **E**. Selected metabolites with different direction of change in lysates and media of mutant iPSC compared to the corrected lines. **F**. Enriched pathways change in lysates and media of mutant iPSC compared to the corrected lines. **G**. qPCR of transporters CAT1, LAT1 and ODC in corrected and mutant iPSC lines. Volcano plot fold change threshold = 2, p-value threshold: 0.01. Statistical significance in B and D based on wilcoxon test.

## Discussion

In this article, we have presented a comprehensive analysis of metabolic changes in the media and lysates from patient-derived iPSC and their corrected counterparts. The genetic alteration results likely results in major changes in the protein structure, affecting it’s secondary and tertiary structure, and likely resulting in protein instability and degradation. The absence or dysfunction of OAT leads to a complex reshaping of the cellular metabolic pathways, which may explain why the dietary changes do not suffice to stop the progression of the disease.

Here, we show intricated metabolic dynamics inside of the cell, as well as their effects on the “circulating” levels of metabolites and the restoration of metabolic homeostasis upon correction of the mutation. Previous research identified the cationic amino acid transporter 1 (CAT1) as possible therapeutic target to regulate ornithine’s effect of retinal cells^19^. Our study shows the decrease of CAT1 expression in corrected cells, indicating a possible upregulation in attempts to keep metabolic homeostasis. We additionally observe an upregulation of ODC in mutant cells, proposing the processing of excessive ornithine towards polyamines. These results support previous hypothesis proposing polyamines excess as a mechanistic feature of the disease ^20–22^. Moreover, eye histology studies have shown focal photoreceptor atrophy around atrophic lesions, and reversible OAT inhibition in human retinal pigment epithelium cells (RPE), as well as non-human primate experiments where ornithine was injected intravitreally showed the destruction of RPE and photoreceptors ^23–25^. Yet, after all these experiments it remains inconclusive to date if hyperornithinaemia affected polyamine pathways and what exactly happens inside the cells. Despite the likely multifactorial nature of the pathogenesis of the disease, the significant increase in intracellular GABA in our mutant cell led us to propose that it is GABA, rather than ornithine or polyamines themselves, one of the central players. With a strong presence in liver and neurons, in one hand it might explain the systemic effects of ornithine accumulation, by a consequent overload of intracellular GABA and an activation of multiple related pathways. On the other hand, GABA itself might be affecting neurons in the eye. In accordance, some epileptic patients treated with vigabatrin present visual field defects^26^. However, GABA has not been widely studied in the context of gyrate atrophy. Hence, deeper metabolomic research, possibly aided by the differentiation of patient-derived iPSC into neurons, hepatocytes, and/or RPE cells, might shed some light into the role of GABA in the tissue specificity of the disease.

## Supplementary Figures

**Figure S1.**
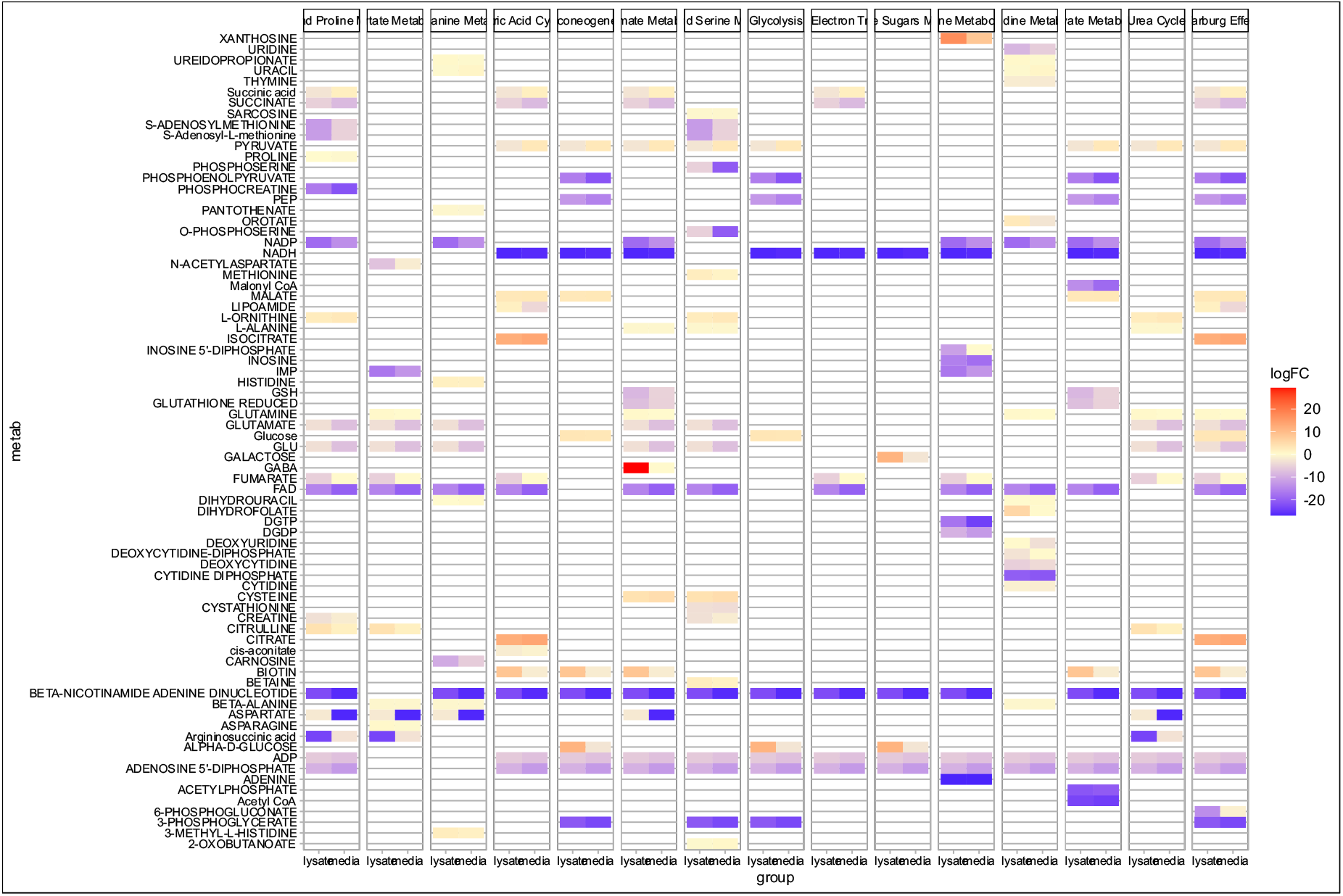
Normalised levels of all changed metabolites in cell lysate and cell media, by their consideration in enriched pathway analysis in Metaboanalyst.

## Materials and methods

### Cell culture

We generated iPSC from skin biopsies from two non-related voluntary female patients, diagnosed with HOGA in their childhood (ethical permit: #HUS/2754/2019). The reprogramming and characterisation of the cells has been reported and described elsewhere^14^.

The patients’ iPSC lines and their genetically corrected counterparts wer cultured on Matrigel (Corning) coated plates in E8 medium (ThermoFisher Scientific) and kept at 37 °C and 5% CO2, with media change every other day. All cell lines tested negative for mycoplasma and present normal karyotype by G-banding (Ambar Anàlisis Mèdiques).

### Metabolomics analysis

Wild type and supplemented TYMS knock out cells were seeded in Matrigel coated 6 well plates and expanded to 70-80% confluency. 6 replicated per genotype were collected and analysed on Thermo Q Exactive Focus Quadrupole Orbitrap mass spectrometer coupled with a Thermo Dionex UltiMate 3000 HPLC system (Thermo Fisher Scientific, Inc.). The HPLC was equipped with a hydrophilic ZIC-pHILIC column (150 × 2.1 536 mm, 5 μm) with a ZIC-pHILIC guard column (20 × 2.1 mm, 5 μm, Merck Sequant). 5 μl of the samples were injected into the LC-MS after quality controls in randomized order having every tenth sample as blank. A linear solvent gradient was applied for separation, in decreasing organic solvent (80–35%, 16 min) at 0.15 ml/min flow rate and 45 °C column oven temperature. The mobile phases included: aqueous 200 mmol/l ammonium bicarbonate solution (pH 9.3, adjusted with 25% ammonium hydroxide), 100% acetonitrile and 100% water. The ammonium bicarbonate solution was kept at 10% throughout the run, resulting in a steady concentration of 20 mmol/l. Metabolites were analysed using a mass spectrometer equipped with a heated electrospray ionization (H-ESI) source using polarity switching and following setting: resolution of 70,000 at m/z of 200; spray voltages: 3400 V for positive and 3000 V for negative mode; sheath gas: 28 arbitrary units (AU), and auxiliary gas: 8 AU; temperature of the vaporizer: 280°C; temperature of the ion transfer tube: 300 °C. The instrument control was conducted with the Xcalibur 4.1.31.9 software (Thermo Scientific). The peaks for metabolites were confirmed with commercial standards (Sigma-Aldrich). Data quality was monitored using a parallel in-house quality control cell line extracted similarly to other samples. The final peak integration was done with the TraceFinder 4.1 SP2 software (Thermo Scientific) and the peak area data was exported as excel file for further analysis. The normalisation of peak area of each metabolite was done with the sum of the absolute peak areas of all metabolites in the same sample. These data were then analysed in R using the limma package^27^, and enrichment analysis was conducted in MetaboAnalyst 6.0. Statistical analysis based on non-parametric Wilcoxon test.

### Protein modelling

*In silico* visualisation and modelling of the OAT protein was conducted in Uniprot (P04181), using its incorporated Alphafold feature. Modelling of the substitution was done in AlphaFold ^28^.

